# Fold-Conditioned De Novo Binder Design via AlphaFold2-Multimer Hallucination

**DOI:** 10.1101/2025.07.02.662497

**Authors:** Khondamir. R. Rustamov, Artyom Y. Baev

## Abstract

*De novo* protein binder design has been revolutionized by deep learning methods, yet controlling binder topology remains a challenge. We introduce a fold-conditioned AlphaFold2-Multimer hallucination framework – FoldCraft – guided by a contact map similarity loss, enabling precise generation of binders with user-defined structural folds. This single loss function enforces fold-specific geometry while implicitly optimizing AlphaFold confidence metrics. We demonstrate the method’s versatility by designing binders with six distinct topologies. Compared to RFdiffusion, FoldCraft yields higher structural and binding confidence. Applied to VHH nanobody design against four therapeutically relevant targets, our method outperforms RFAntibody in AlphaFold3-based evaluations. FoldCraft offers a general, efficient strategy for structure-guided binder design, expanding the accessible fold space for protein engineering and enabling robust nanobody generation with improved success rates.

## Introduction

Proteins are the fundamental macromolecules responsible for executing nearly all cellular functions. A substantial portion of these functions arises through protein–protein interactions (PPIs), which underpin key biological processes such as signal transduction, immune recognition, and structural organization. The ability to design synthetic protein binders that selectively modulate PPIs has transformative potential in therapeutic development, diagnostics, and synthetic biology [1-3]. However, conventional protein design approaches remain prohibitively slow, labor-intensive, and resource-demanding.

Recent advances in deep learning–based protein structure prediction have catalyzed a paradigm shift in biomolecular modeling. Tools such as AlphaFold2 (AF2) [4] and RoseTTAFold2 (RF2) [5] have demonstrated unprecedented accuracy in predicting protein structures and modeling PPIs, laying the foundation for structure-guided binder design [6]. These developments have enabled new classes of deep learning–driven generative models, including RFdiffusion [7], which hallucinates protein backbones, and ProteinMPNN [8], which designs sequences for predefined scaffolds. RFdiffusion-AA [5] further extends this framework to generate binders against a broader range of biomolecular targets, including nucleic acids and small molecules. Despite these advances, diffusion-based methods often require the generation of thousands of candidates followed by extensive in vitro screening to identify functional binders – limiting their overall efficiency.

Alternative strategies such as BindCraft [9], which employs backpropagation through AF2-Multimer combined with SolMPNN [10], for sequence optimization, have shown improved success rates. Likewise, BoltzDesign1 [11] based on inversion of the open-source Boltz1 structure prediction model [12], has demonstrated enhanced designability across a range of molecular targets, surpassing RFdiffusion-AA in computational metrics.

Despite these advances, most state-of-the-art methods are constrained to binders with regular secondary structures and struggle to generate designs with predefined complex topologies, such as nanobody and antibody folds. A recent study [13] attempted to address this by fine-tuning RFdiffusion-AA and RoseTTAFold for de novo antibody design, but the method required experimental screening of nearly 9,000 designs to identify nanomolar binders, highlighting the limitations in efficiency and fold control.

Here, we introduce FoldCraft – a fold-conditioned binder design strategy based on AlphaFold2-Multimer hallucination. By incorporating contact map similarity loss, our approach enables the generation of binder sequences that conform to user-specified topologies while forming high-confidence interfaces with target proteins. We apply this framework to the de novo design of structurally diverse binders, including immunoglobulin-like and nanobody folds, and benchmark it against leading generative models.

## Methods

### Contact probability maps (cmaps) similarity loss

To guide binder design toward a specified structural fold, we introduce a loss function based on the similarity of contact probability maps (cmaps) between a reference structure and the predicted binder. First, a template cmap is constructed from a reference protein exhibiting the desired fold. These reference structures were generated using AlphaFold2 predictions based on masked experimental structures (see Table S1). Intra-chain contact probabilities were computed from the AlphaFold2 distogram output, using a 14 Å threshold to define contacts within the monomeric template structure [4, 14].

This monomeric cmap was then adapted for the binder–target complex by masking all intra-chain contacts in the target and defining desired inter-chain contacts between known interface residues of the target and the binder. These defined inter-chain contacts help guide the design toward both structural fidelity and target engagement.

During each hallucination step, AlphaFold2-Multimer predicts the 3D structure of the binder–target complex. From the predicted distogram, we compute a new cmap using a 21.7 Å cutoff for inter-chain contacts and 14 Å for intra-chain contacts. The difference between the hallucinated and template cmaps is used to compute the loss, which is used to optimize the binder sequence toward the desired fold while promoting productive binding with the target.

### Binder design protocol

We employed the ColabDesign implementation of AlphaFold2-Multimer to perform hallucination-based binder sequence and structure co-generation. The design process begins with initializing a random amino acid sequence for the binder. AlphaFold2-Multimer then predicts the structure of the binder–target complex in a single-sequence mode for the binder, while the target protein is provided as a structural template [15, 16]. From the predicted distogram, a contact probability map (cmap) is computed and compared to the template cmap to calculate the similarity loss. This loss is then backpropagated through the AlphaFold2-Multimer network, and the binder sequence is optimized using stochastic gradient descent to iteratively minimize the cmap similarity loss [17].

We adopted the three-stage sequence optimization scheme from ColabDesign (100 steps – logits, 100 steps – softmax, 20 steps – one hot encoding) to iteratively optimize binder sequence [9]. Once binder sequences were generated, we further refined them using ProteinMPNN or SolMPNN to enhance predicted stability, solubility, and expression [8, 18].

In our default pipeline, SolMPNN was used to optimize residues outside a 4 Å radius from the binder–target interface, preserving the interface geometry while improving biophysical properties. To evaluate alternative sequence optimization strategies, we also tested full-sequence redesign using both ProteinMPNN and SolMPNN. For all optimization protocols, we used a temperature of 0.1 and set backbone noise to 0.0.

### In silico benchmarking

To evaluate the performance of FoldCraft, we benchmarked it against SOTA models across two distinct design tasks: generation of binders with diverse structural folds and design of VHH nanobody binders.

We compared our method with RFdiffusion (run using default settings) for designing binders with six diverse fold topologies against the PD-L1 receptor. The receptor structure was obtained from AlphaFold protein structure database [19]. For each selected fold, we generated 10 binder backbones using either RFdiffusion or FoldCraft. Each backbone was then used to generate 10 sequences using ProteinMPNN or SolMPNN. Finally, all sequences were subsequently re-evaluated using AlphaFold2-ptm (3 recycles, 2 models). Binder sequences were predicted in single-sequence mode, while the target protein structure was provided as a fixed structural template.

To assess FoldCraft’s ability to design VHH nanobody folds de novo, we benchmarked it against RFAntibody [13]. RFAntibody was run locally using default settings. For each of four targets (PD-1, PD-L1, IFNAR2, and EGFR), we generated 40 nanobody structures using the default VHH framework, with CDR loops masked in both protocols. For RFAntibody, sequence optimization was performed using the default ProteinMPNN-based CDR loop redesign, as described in the original publication. In our protocol, ProteinMPNN or SolMPNN was used to optimize non-interface regions of the hallucinated binder sequences. For each nanobody structure, five sequences were generated using either ProteinMPNN or SolMPNN.

AlphaFold3 (AF3) was employed as the oracle model for evaluating nanobody-target complexes, as it has been shown to outperform AlphaFold2-Multimer and antibody-finetuned RoseTTAFold in predicting antibody–antigen interactions [13, 20, 21]. AF3 was run locally: target structures were provided via structural templates, while VHH binders were predicted without MSA but with template search enabled. All predictions were generated using one diffusion sample. Binder designs were evaluated based on three key AlphaFold3 metrics: default ipTM score, the average plDDT score of Cα atoms within the binder, and the mean predicted aligned error for inter-chain residue-residue pairs between binder and target (iPAE) [22-24].

## Results

### Fold conditioned de novo design of protein binders

Recent advances in *de novo* protein binder design have been significantly accelerated by deep learning–based algorithms such as RFdiffusion and BindCraft [7, 9]. In this work, we aimed to extend the binder design framework by introducing fold-conditioning, enabling the design of proteins with predefined structural topologies.

The first stage of the BindCraft pipeline utilizes ColabDesign to hallucinate binder sequences and structures via backpropagation through the AlphaFold-Multimer model. ColabDesign supports multiple binder design strategies, including *de novo* generation and redesign of existing binder templates using a range of loss functions. While the original BindCraft study primarily focused on designing binders with regular secondary structures (i.e., varying proportions of α-helices and β-sheets), it did not explicitly enforce structural motifs such as antibody folds. This dependence limits their ability to generate binders with predefined scaffolds or folds *de novo*.

To overcome this limitation, we drew inspiration from RFAntibody, which conditions designs on antibody-like folds using structural templates that encode pairwise distances and dihedral angles characteristic of canonical antibody or nanobody frameworks [13]. Analogously we adapted the ColabDesign binder design protocol to similarly bias designs toward conditioned folds.

Our fold-conditioned binder design pipeline uses ColabDesign implementation of AF2-Multimer to optimize binder sequences by minimizing the difference of binder structure to desired fold and maximizing its contacts with target through iterative backpropagation. AF2-Multimer provides pairwise distance distributions, from which we compute contact probability maps (cmaps). These maps quantify the likelihood that a given residue pair is within a specified distance threshold (14 Å for intra-chain contacts, 21.7 Å for inter-chain contacts). Specifically, we defined the desired binder fold via a template cmap (calculated from a monomeric reference structure), and masked all contacts within the target protein, as well as inter-chain contacts not involved in binding. To promote target engagement, we explicitly defined inter-chain contacts between selected target hotspots and corresponding binder residues (Fig. 2A). To guide the AF2-Multimer hallucination process toward generating binder structures that adopt a desired fold, we define a loss based on the similarity of contact maps between the predicted and target (conditioned) structures. This loss is computed as a Root Mean Square Error (RMSE) over a masked subset of residue–residue contact pairs:

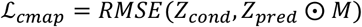

During design, a random initial binder sequence is generated, and the binder–target complex is predicted using AlphaFold2-Multimer. The cmap of predicted complex structure (*Z_pred_*) is compared to the predefined template (*Z_cond_*) and binary mask *M* is applied to restrict the comparison to relevant contact regions. The loss is computed as *RMSE* between masked predicted and predefined cmaps and the initial binder sequence is optimized to reduce this loss. This process is iterated until the binder adopts the desired structural fold and forms the intended interface with the target.

As previous studies have shown that sequences derived from hallucination protocols often exhibit poor stability and expression in experimental settings, we incorporated ProteinMPNN and SolMPNN [8, 10] to refine our final designs. Specifically, residues located outside of the predicted binding interface were redesigned to improve predicted solubility, stability, and expression potential.

Finally, all optimized binder sequences were structurally predicted in complex with the target using AlphaFold2. These final complexes were evaluated using AlphaFold confidence metrics (pLDDT, ipAE), and assessed for structural consistency with the intended fold template to validate the success of fold conditioning.

### Designing binders with diverse folds

To evaluate the ability of FoldCraft to guide binder design toward specified structural topologies, we selected six representative protein folds with diverse architectures: Top7, the first *de novo* designed protein [17, 25], β-barrel, Ig-like fold, TIM barrel – more complex folds which have been succesfully generated with AF2 hallucination protocols and SolMPNN [10] β-solenoid and Ankyrin repeat, naturally occurring representatives of solenoid folds, which has been recently explored using custom loss functions in ColabDesign [26]. For each fold, we used FoldCraft to design fold-conditioned binders targeting PD-L1, a therapeutically relevant immune checkpoint receptor for which several *de novo* binders have been previously validated [9, 27, 28]. In all six cases, we successfully generated binders adopting the intended structural fold, with 8–45% of the designs achieving an RMSD of less than 3.5 Å relative to the monomeric fold template (Fig. 3A).

We next sought to evaluate whether our contact map similarity loss implicitly optimizes the loss functions used in BindCraft, which employs six AF2-Multimer–based metrics during binder hallucination: plDDT, ipTM, PAE, iPAE, intra-chain contacts within binder (con), and inter-chain contacts (i_con). In our experiments, we observed strong correlations between cmap similarity loss and intra-chain metrics such as plDDT, PAE, and con, and moderate correlations with inter-chain metrics ipTM, iPAE, and i_con (Fig. 2C). These results indicate that our contact map loss not only steers designs toward the desired fold but also implicitly improves AlphaFold confidence metrics that assess both the quality of the binder structure and its predicted interaction with the target protein.

We then benchmarked our pipeline against RFdiffusion with fold-conditioning across the same six structural folds. In addition, we compared two sequence optimization strategies for post-design refinement: (i) full-sequence optimization using ProteinMPNN or SolMPNN, and (ii) optimization restricted to non-interface regions, as implemented in BindCraft. All final designs were predicted using AlphaFold2, and successful binders were identified using established thresholds based on AlphaFold confidence metrics (Fig. 3C).

Our results show that FoldCraft consistently outperforms RFdiffusion in generating binders that adopt the desired structural folds. While RFdiffusion followed by ProteinMPNN produced designs with high plDDT scores in some cases, the overall binding confidence (e.g., ipTM and iPAE) and fold accuracy (RMSD to the template structure) were substantially lower compared to our method (Fig 3B).

Finally, we compared the effects of full-sequence versus non-interface–restricted optimization on design quality. Full-sequence optimization using ProteinMPNN or SolMPNN generally yielded higher plDDT scores, indicating improved structural confidence. However, designs with non-interface regions optimized only – achieved superior binding scores (ipTM and iPAE) and lower RMSD to the fold template, suggesting that restricting optimization to non-interface regions can better preserve fold-specific geometries while maintaining favorable binding predictions (Fig. 3B).

### De novo design of VHH single domain nanobodies

The primary goal of this study was the *de novo* design of VHH nanobody binders using FoldCraft. Antibody and nanobody folds have long represented a major challenge for deep learning-based *de novo* protein design methods [29-31]. While this challenge has recently been addressed by the RFAntibody model – which applies template-guided diffusion to generate diverse antibody-like folds – the overall success rate of the method remains extremely low, requiring large-scale experimental screening to identify functional binders [13].

To evaluate the effectiveness of our strategy, we designed VHH fold-conditioned binders against four therapeutically relevant cell surface receptors: PD-1, PD-L1, IFNR2, and EGFR, each previously validated as successful binder design targets [9, 27]. Structures for all targets were obtained from AlphaFold structure prediction database [19]. Given the structural flexibility of the CDR loops in VHH domains, we masked these regions when generating our fold-conditioning contact maps (Fig. 1C).

**Figure 1.**
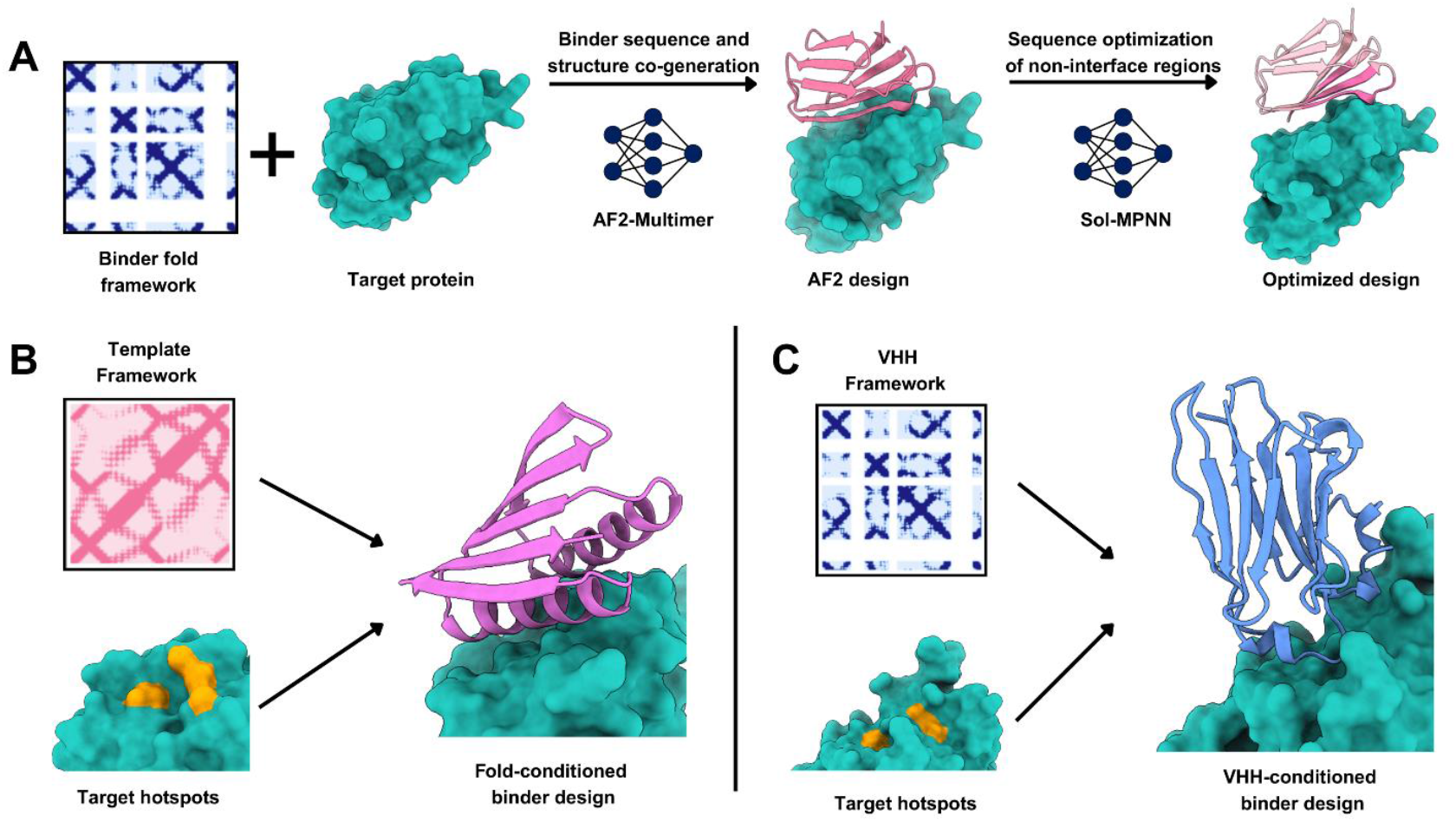
Fold-conditioned *de novo* binder design pipeline. **A)** Overview of the design protocol. Target protein structure is used as inputs to guide binder sequence and structure co-generation using AlphaFold2-Multimer (AF2-Multimer) based on cmap similarity loss to predefined fold-conditioned template cmap. The hallucinated complex is then optimized by sequence optimization of non-interface regions using SolMPNN to improve solubility and stability. **B)** Fold conditioning using a template structural framework. The template cmap defines intra-chain structural constraints, while specific inter-chain interactions are biased toward predefined target surface hotspots. **C)** VHH nanobody design using a VHH-specific template.

**Figure 2.**
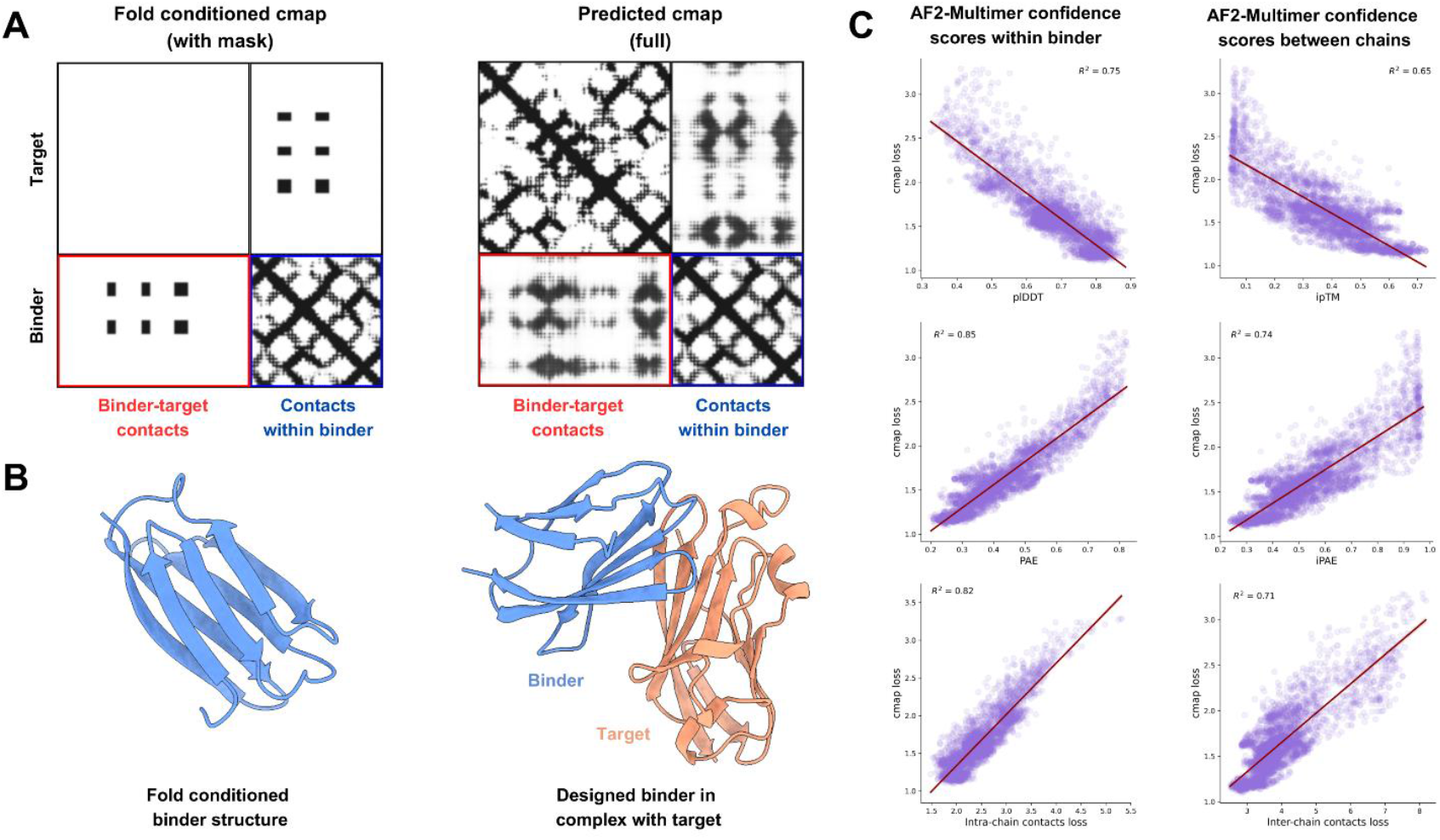
Contact map conditioning and its correlation with AlphaFold2-Multimer confidence metrics. **A)** Schematic illustration of fold-conditioned binder design using cmap similarity loss. The fold-conditioned input cmap (left) includes masked intra-chain contacts of the target, explicit intra-chain contacts within the binder (blue), and specified inter-chain contacts between target hotspots and the binder (red). The predicted contact map (right) is generated during AlphaFold2-Multimer hallucination and used to compute the root mean square error (RMSE) against the fold-conditioned input. **B)** Template structure for computing the fold-conditioned cmap (left) and designed binder in complex with target (right).**C)** Correlation between cmap similarity loss and AF2-Multimer confidence scores.

**Figure 3.**
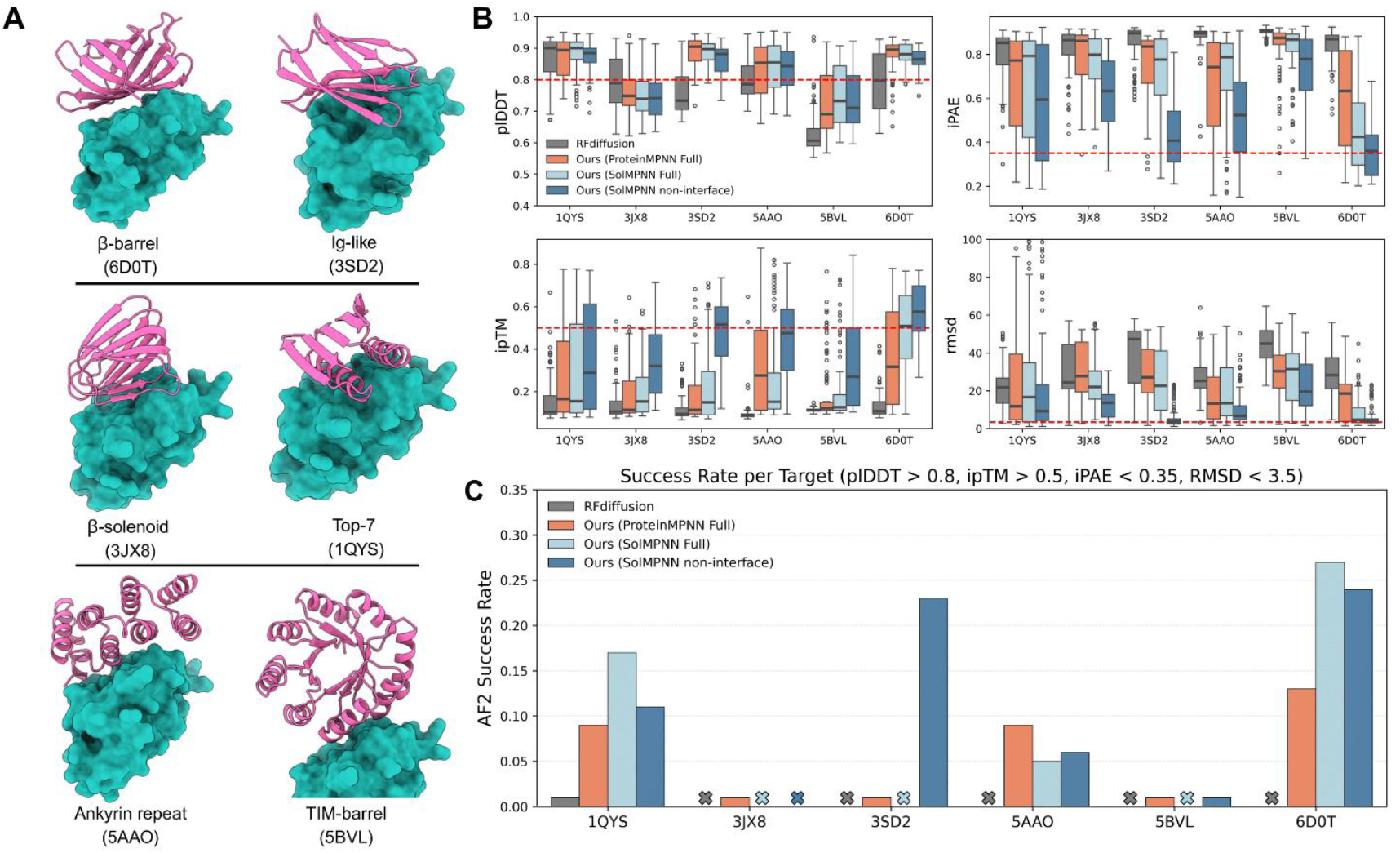
Benchmarking fold-conditioned binder design across diverse protein folds. **A)** AF2 predicted structures of designed binders with six distinct target folds used for conditioning. All binders were generated using FoldCraft and shown bound to their respective target protein (PD-L1). **B)** Comparison of AF2-Multimer prediction metrics—plDDT, ipTM, iPAE, and RMSD to template fold – across different optimization protocols. Dashed red lines indicate recommended thresholds for high-confidence predictions. **C)** AF2 *in silico* success rates per fold type, defined as the percentage of designs satisfying plDDT > 0.8, ipTM > 0.5, iPAE < 0.35, and RMSD < 3.5 Å.

We benchmarked our fold-conditioned approach against the RFAntibody model followed by sequence optimization using ProteinMPNN. For each target, we generated 40 backbone designs, and for each backbone, five sequence variants were created using either SolMPNN or ProteinMPNN.

All resulting binder–target complexes were structurally predicted using the AlphaFold3 model [20], which has been shown to outperform AlphaFold2 and antibody-finetuned RoseTTAFold models in predicting antibody–antigen interactions [13, 21]. Target structures were predicted using input structural templates, while VHH binders were predicted with template search, but without MSA, as described in RFAntibody paper.

We assessed the predicted complexes using three AlphaFold3-based criteria:

1. plDDT > 70 and ipAE < 10 (or <15 for more relaxed filters), as adopted in BoltzDesign1 [11, 22, 32],
2. ipTM threshold, which has been reported to correlate with accurate antibody–antigen binding [9, 13, 21]

As shown in Figure 4A, FoldCraft successfully generated VHH binders across all four targets. While the RFAntibody + ProteinMPNN pipeline produced nanobody designs with high plDDT scores, none of these designs passed the AlphaFold3 interface filters, indicating poor predicted binding (Fig. 4B). In contrast, our method achieved 0.5–19.5% in silico success rates, demonstrating the effectiveness of FoldCraft in generating structurally and functionally plausible VHH binders.

**Figure 4.**
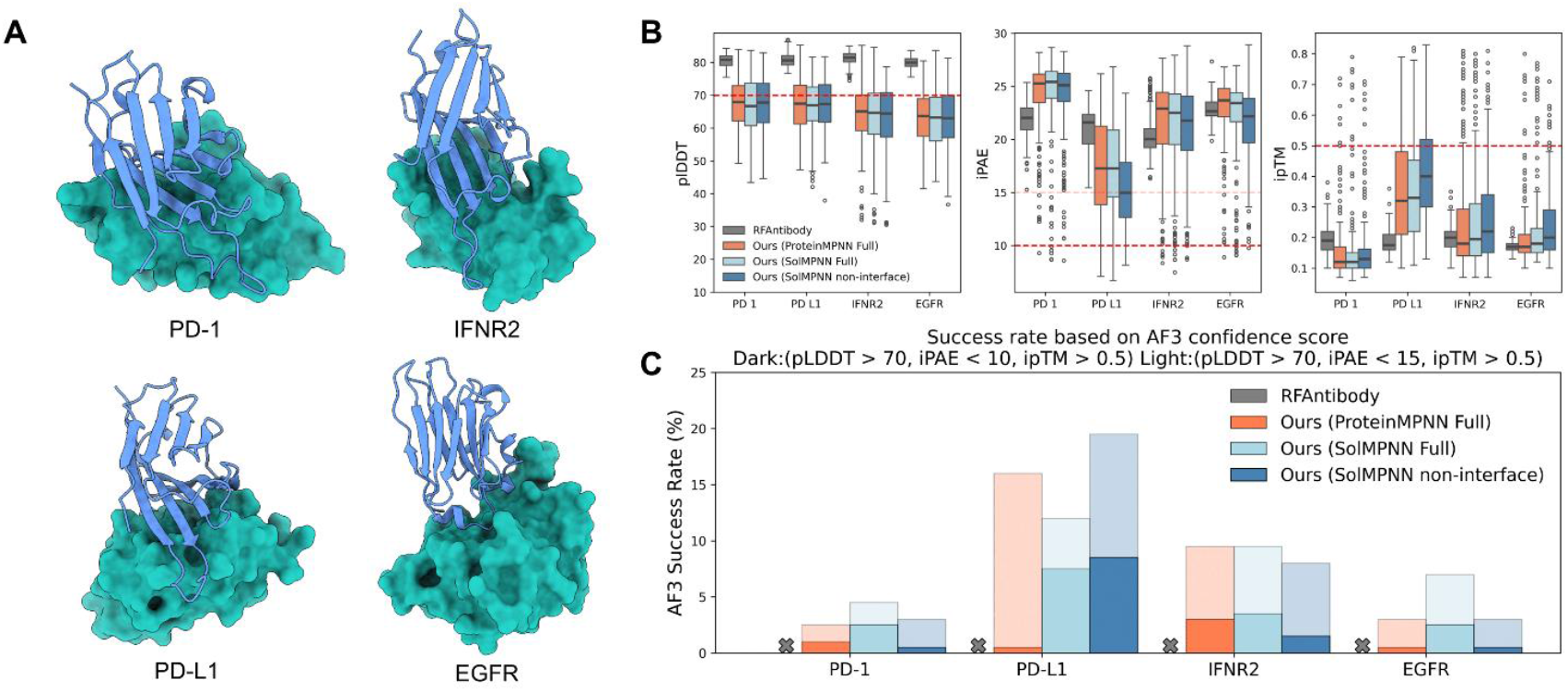
*De novo* design of VHH nanobody binders FoldCraft. **A)** AF3 predicted complexes of fold-conditioned VHH binders (blue) targeting four therapeutically relevant receptors: PD-1, PD-L1, IFNR2, and EGFR (cyan). **B)** Comparison of AF3 confidence metrics for nanobody binders designed with FoldCraft versus RFAntibody + ProteinMPNN pipeline. Dashed red lines denote thresholds for high-confidence binder predictions. **C)** *In silico* success rates for both methods, defined by AF3 confidence scores. Dark bars correspond to stricter criteria (plDDT > 0.7 and iPAE < 10), while light bars reflect relaxed thresholds (plDDT > 0.7 and iPAE < 15).

Previous studies have shown that ProteinMPNN struggles to recover sequences compatible with antibody structures, which may underlie the poor binding confidence observed with RFAntibody-designed sequences. To evaluate whether ProteinMPNN could more effectively optimize single-domain antibody sequences when combined with our method, we applied ProteinMPNN and SolMPNN to redesign the full sequence of FoldCraft-generated VHH binders. Notably, both optimization strategies improved predicted binding metrics, resulting in higher in silico success rates compared to RFAntibody (Fig. 4B).

Upon structural analysis of the top-performing designs, we observed that many predicted nanobody binders engaged in side-docking interactions, a phenomenon previously reported in the RFAntibody study. This recurring artifact may arise from a bias in the training datasets of AlphaFold3 and RoseTTAFold, where the majority of nanobody–antigen complexes exhibit side-dock geometries [13].

Overall, FoldCraft significantly improves the designability of VHH scaffolds compared to the RFAntibody framework, enabling more consistent generation of high-confidence *de novo* nanobody binders.

## Conclusion

In this study, we present a generalizable, fold-conditioned de novo binder design framework – FoldCraft – that leverages AF2-Multimer hallucination guided by a contact map similarity loss. Unlike existing methods that require elaborate multi-loss strategies or diffusion-based backbone generation, FoldCraft simplifies the optimization process by enforcing structural fidelity through a single loss function derived from contact map similarity [9]. This innovation allows our pipeline to condition designs on specific structural folds while implicitly optimizing AlphaFold confidence metrics, including pLDDT, ipTM, and iPAE.

We first evaluated the versatility of FoldCraft by designing binders adopting six distinct protein topologies, ranging from canonical folds like Top7 and β-barrels to noncanonical architectures such as β-solenoids and TIM barrels. Across all fold types, FoldCraft consistently outperformed RFdiffusion – currently considered a state-of-the-art method for de novo binder design – both in terms of structural fidelity and AlphaFold confidence metrics. Importantly, we demonstrate that our single-loss strategy can capture and enforce complex structural constraints without additional auxiliary terms, making it more robust and broadly applicable across diverse protein families.

We further applied FoldCraft to the de novo design of VHH nanobody binders, a class of small, single-domain antibodies that pose significant challenges for generative models due to their complex fold topology and conformationally flexible loops. While RFAntibody recently addressed this challenge through template-guided diffusion, the approach requires large-scale screening of thousands of designs to identify functional candidates [13]. In contrast, FoldCraft generated high-confidence VHH binders against four therapeutically relevant targets (PD-1, PD-L1, IFNR2, and EGFR), achieving in silico success rates of up to 19.5% as predicted by AlphaFold3. Notably, these binders consistently passed multiple structural filters (pLDDT > 70, ipAE < 10, ipTM > 0.5), and none of the RFAntibody-designed binders met these thresholds.

An additional strength of FoldCraft lies in its compatibility with post-hallucination sequence optimization protocols. We observed that full-sequence redesign with ProteinMPNN or SolMPNN improves overall structural confidence (plDDT), while interface-restricted optimization better preserves binding geometry (lower ipAE and RMSD to the fold template). This flexibility shows high designability of FoldCraft generated designs and enables users to tailor optimization strategies depending on their desired balance between stability, solubility, and binding specificity. Despite these advantages, certain limitations remain. One recurring issue was the tendency of the oracle model (AlphaFold3 or RoseTTAFold) to predict side-docking interactions for VHH designs – likely reflecting biases in the training data, which overrepresent such binding modes. Addressing this artifact may require improvements in the training datasets or fine-tuning models to support more natural antibody docking geometries [13].

Additionally, FoldCraft did not generalize to scFv antibody topologies, unlike the RFAntibody model. This limitation likely stems from the lower accuracy of AF2-Multimer in docking complex antibody folds to their targets. Advances in structure prediction models used within hallucination workflows may help overcome this challenge. In particular, incorporating open-source reproductions of AlphaFold3, such as Boltz (as used in the BoltzDesign1 pipeline), holds strong potential for enabling accurate scFv antibody binder design [11, 12, 20].

Moving forward, future work will focus on expanding FoldCraft’s applicability to more complex antibody scaffolds and experimentally validating computationally designed binders.

In conclusion, FoldCraft offers a powerful and general-purpose platform for structure-guided de novo binder design. It achieves strong performance across diverse folds, enables efficient nanobody generation, and surpasses state-of-the-art models in in silico success. This work opens the door for more targeted and reliable de novo antibody and binder design pipelines, with promising applications in therapeutic development, diagnostics, and synthetic biology.

## Code availability

The full code for FoldCraft is available on GitHub (https://github.com/KhondamirRustamov/FoldCraft).

## Supplementary Materials

**Supplementary table 1.**
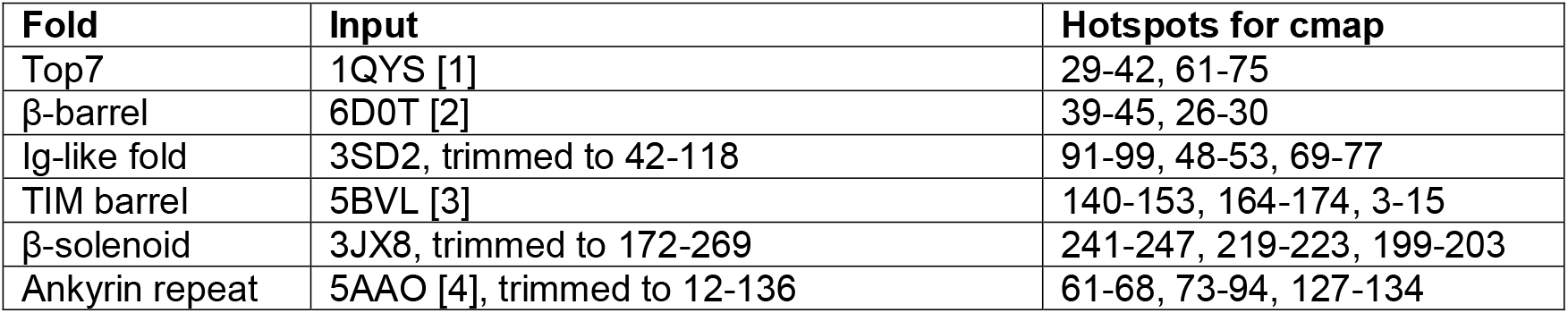
Binder template structures for fold-conditioned design.

**Supplementary table 2.**
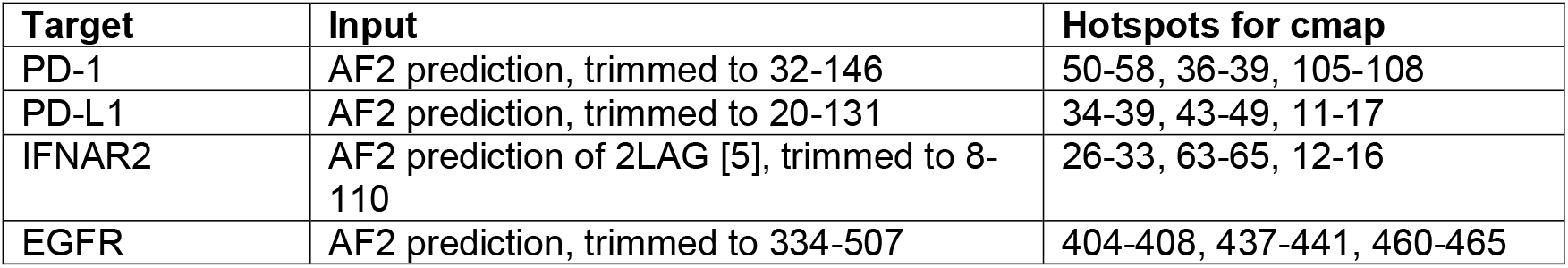
Target structural templates for binder design.

