## Supplemental Table 1 for "Fold-Conditioned De Novo Binder Design via AlphaFold2-Multimer Hallucination"

**Supplementary Materials**

**Supplementary table 1.** Binder template structures for fold-conditioned design.

| **Fold** | **Input** | **Hotspots for cmap** |
| --- | --- | --- |
| Top7 | 1QYS [1] | 29-42, 61-75 |
| β-barrel | 6D0T [2] | 39-45, 26-30 |
| Ig-like fold | 3SD2, trimmed to 42-118 | 91-99, 48-53, 69-77 |
| TIM barrel | 5BVL [3] | 140-153, 164-174, 3-15 |
| β-solenoid | 3JX8, trimmed to 172-269 | 241-247, 219-223, 199-203 |
| Ankyrin repeat | 5AAO [4], trimmed to 12-136 | 61-68, 73-94, 127-134 |

**Supplementary table 2.** Target structural templates for binder design.

| **Target** | **Input** | **Hotspots for cmap** |
| --- | --- | --- |
| PD-1 | AF2 prediction, trimmed to 32-146 | 50-58, 36-39, 105-108 |
| PD-L1 | AF2 prediction, trimmed to 20-131 | 34-39, 43-49, 11-17 |
| IFNAR2 | AF2 prediction of 2LAG [5], trimmed to 8-110 | 26-33, 63-65, 12-16 |
| EGFR | AF2 prediction, trimmed to 334-507 | 404-408, 437-441, 460-465 |

**References.**
